# Continuous detection of glucose and insulin in live animals

**DOI:** 10.1101/2020.01.22.916106

**Authors:** Mahla Poudineh, Caitlin L. Maikawa, Eric Yue Ma, Jing Pan, Dan Mamerow, Yan Hang, Sam W. Baker, Ahmad Beirami, Michael Eisenstein, Seung Kim, Jelena Vučković, Eric A. Appel, H. Tom Soh

## Abstract

Real-time biosensors that can continuously measure circulating biomolecules *in vivo* would provide valuable insights into a patients’ health status and their response to therapeutics even when there is considerable variability in pharmacokinetics and pharmacodynamics across patient populations. Unfortunately, current real-time biosensors are limited to a handful of analytes (*e.g.* glucose and blood oxygen) and are limited in sensitivity (high nanomolar). In this work, we describe a general approach for continuously and simultaneously measuring multiple analytes with picomolar sensitivity and sub-second temporal resolution. As exemplars, we report the simultaneous detection of glucose and insulin at picomolar concentrations in live diabetic rats. Using our system, we demonstrate the capacity to resolve inter-individual differences in the pharmacokinetic responses to insulin and discriminate profiles from different insulin formulations at a high temporal resolution. Critically, our approach is general and could be readily modified to continuously and simultaneously measure other circulating analytes *in vivo* by swapping the affinity reagents, thus making it a versatile tool for biomedical research.

## Introduction

Technologies that can continuously measure circulating biomolecules *in vivo* would have a transformative impact towards the vision of precision medicine^1^. Such tools could provide valuable insights into a patients’ health status and their response to therapeutics, allowing clinicians to tailor therapeutic regimens to consistently deliver maximum efficacy with minimal side-effects even when there is considerable variability in pharmacokinetics and pharmacodynamics across patient populations. Early efforts toward this end have already had a considerable impact; for example, continuous glucose monitors have significantly reduced the burden for diabetes patients and increased the time where patients’ blood glucose levels are within the euglycemic range^2^. Similar monitoring could prove invaluable for managing a host of other disease states^3–6^. Unfortunately, it remains exceedingly difficult to achieve continuous *in vivo* detection of biomolecules, and to date, continuous real-time sensors have been limited only to a handful of analytes such as glucose^7^, lactate^8^, and blood oxygen^9^.

Recently, a new generation of biosensors has emerged that can continuously measure other types of biomolecules *in vivo*. For example, our group demonstrated the first platform for the continuous detection of small-molecule drugs in the bloodstream of live animals using an electrochemical sensor based on structure-switching aptamer probes^10^. Using a similar strategy, Arroyo-Currás *et al.* have shown multi-hour monitoring of four drugs (Doxorucibin, Kanamycin, Gentamicin, and Tobramycin) in blood^11^. These platforms represent notable advances in terms of demonstrating the feasibility of continuous *in vivo* molecular detection, but such approaches have been limited in sensitivity. These early biosensors can only measure target molecules at relatively high concentrations (typically in the high nanomolar to micromolar range), and could not measure other clinically important analytes that exist at lower concentrations, for example, in the picomolar range^12^.

Insulin is one of the most important therapeutic molecules, with over 50 million diabetes patients worldwide requiring insulin replacement therapy^13–15^. Accurate insulin dosing is critical, because excess insulin can result in acute hypoglycemia, which can lead to dangerously low blood sugar levels that can lead to death^16,17^. One study from 2015 estimated that consequences of insulin administration-related errors by patients account for more than 97,000 hospital visits annually in the US^18^. Furthermore, it is well-known that patients have diverse responses to insulin, and that insulin absorption rates and clearance rates can differ dramatically among patients—or even in the same patient, depending on factors such as temperature, preparation technique and injection site^19–21^. However, measuring insulin kinetics for individual patients requires frequent blood draws and laboratory analysis which is not practically feasible. As a result, current treatment regimens are planned based on modeled population data and include an element of trial and error due to the abovementioned inter-individual variability. Thus, the capability to continuously measure *in vivo* insulin and glucose concentrations would be highly valuable, enabling clinicians to create optimal therapeutic regimens that are tailored for individual diabetic patients.

To this end, we report a novel system that can continuously and simultaneously measure physiological levels of circulating glucose and insulin *in vivo* with picomolar sensitivity and sub-second temporal resolution. Our assay (called “real-time ELISA” or RT-ELISA) integrates aptamer- and antibody-based molecular probes into a bead-based fluorescence assay, wherein analyte concentrations are measured with a highly sensitive optical readout using a specially designed microfluidic chip. For the first time, we demonstrate the capability to simultaneously measure clinically relevant concentrations of both insulin and glucose in the blood of live diabetic rats. Importantly, we demonstrate the capacity to clearly discriminate inter-individual differences in the pharmacokinetic and pharmacodynamic responses to insulin. Our system is also able to distinguish the clear pharmacokinetic differences between different formulations of insulin, which were specifically developed for short- and long-acting time profiles.

## Results & Discussion

### RT-ELISA detection strategy

RT-ELISA employs two different bead-based target capture strategies to detect both insulin and glucose in parallel. For glucose, we used aptamer probes and a strand displacement strategy^22–24^. Briefly, we hybridized Cy5 fluorophore-conjugated glucose aptamers^25^ with a DNA competitor strand that has been conjugated to a quencher (BHQ2) molecule and coupled these complexes to polystyrene microbeads. The aptamer-DNA competitor complex keeps the fluorophore and quencher in close proximity, producing no signal in the absence of target. When the aptamer binds to glucose, the competitor strand dissociates and alleviates quenching of the fluorophore, producing a signal (Figure 1A, left). For insulin, we developed a fluorescence-based sandwich immunoassay in which microbeads were functionalized with anti-human insulin antibodies as capture reagents, with detection achieved with a second anti-insulin antibody labeled with R-phycoerythrin (R-PE) (Figure 1B, left).

**Figure 1.**
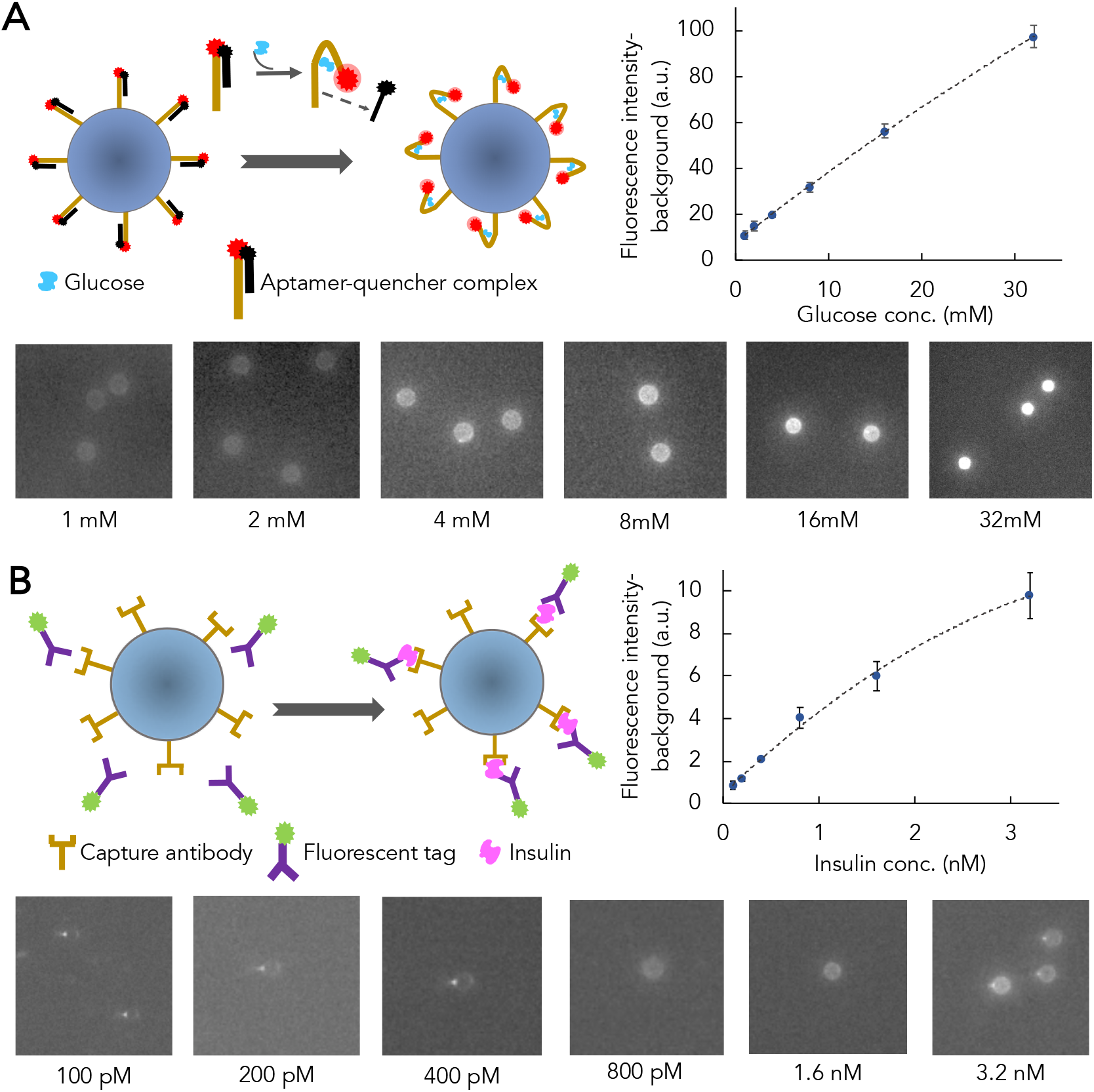
Overview and validation of RT-ELISA assay strategy. RT-ELISA employs two parallel assays to track transient changes in glucose and insulin levels. **A)** 15 μm microbeads are functionalized with a glucose aptamer conjugated with Cy5 fluorophore, hybridized to a DNA competitor conjugated with a quencher (BHQ2). In the presence of glucose, the competitor strand is displaced, eliminating quenching and producing a fluorescent signal (left). For glucose probe validation, microbeads functionalized with the aptamer-competitor complex are incubated with different concentrations of glucose in buffer. After an hour, the beads are washed three times and imaged with a red laser to excite the Cy5 fluorophore (bottom). Plot shows fluorescence signal intensity at each concentration (right). **B)** Insulin is detected in a sandwich assay, with microbead-conjugated capture antibodies and R-phycoerythrin (R-PE)-tagged detection antibodies (left). For insulin probe validation, microbeads functionalized with the capture antibodies are incubated with different concentrations of insulin and R-PE-labeled detection antibodies. After an hour, beads are washed three times and imaged with a green laser to excite the R-PE fluorophore (bottom). Plot shows fluorescence signal intensity at each concentration (right). For each concentration in **A** and **B**, at least 10 beads were measured, and the error bars show standard deviation among the beads. Non-linear regression analysis is used for curve-fitting.

We initially validated the performance of our glucose and insulin assays in buffer. After functionalizing microbeads with the aptamer-DNA competitor complex for glucose detection or insulin capture antibodies, we incubated the beads with different concentrations of glucose (Figure 1A) or insulin plus detection antibodies (Figure 1B). After an hour, we were able to detect bead-target complexes under a fluorescence microscope. We estimated the limit of detection (LOD) of our assay to be three times the standard deviation of the fluorescence signal intensity from a blank sample (see Methods). We achieved a LOD of 3.2 mM and 170 pM for glucose and insulin, respectively. For non-diabetic patients, secreted insulin concentrations are typically ~40 pM when fasting, rising to over 500 pM after meals^26,27^. Peak insulin concentrations in patients with diabetes can be more variable; in patients with type 1 diabetes, typical peak insulin concentrations are dependent on the dose after a prandial bolus are between 200-600 pM^28,29^. Glucose levels rarely deviate from between 3.5-8.0 mM in non-diabetic patients, but in diabetics, concentrations can rise to >20 mM after meals if insulin is not given^27,30^. Based on these metrics, our assay can achieve sufficient sensitivity for measuring physiologically relevant concentrations of glucose and insulin in diabetic patients.

### RT-ELISA device testing and optimization

The RT-ELISA device integrates three modules to achieve continuous, real-time monitoring: (1) a mixing module, which combines molecular probes for analyte detection with the whole blood sample; (2) a depletion module, which minimizes background by reducing the number of blood cells in the sample; and (3) a detection module, which brings the target to the detection window to quantitatively measure its abundance (Figure 2).

**Figure 2.**
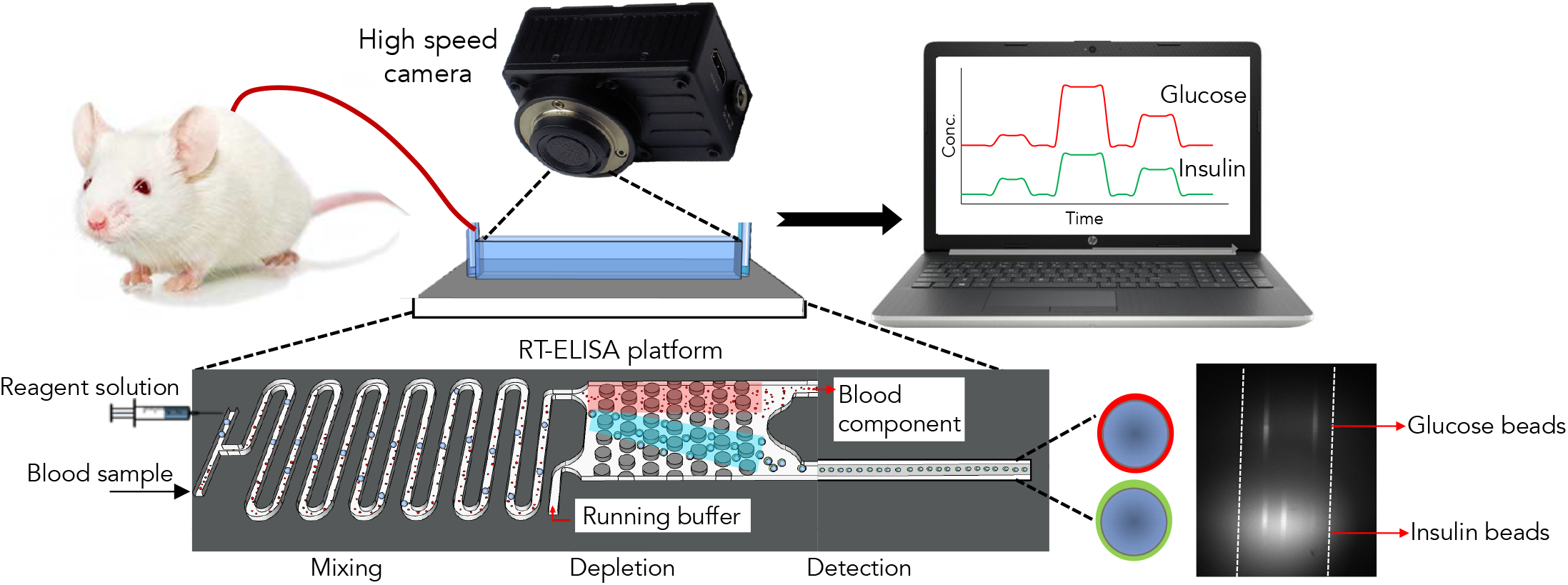
Overview of RT-ELISA platform. The biosensor is connected to a rat through an angio-catheter, with blood injected into the device using a peristaltic pump. The device consists of three modules: a mixing module where blood is combined with detection reagents, a depletion module for eliminating excess blood cells, and a detection module that transfers the fluorescently labeled beads to the detection window.

We used a standard microfluidic device fabrication protocol with glass substrates and polydimethylsiloxane (PDMS) to build the RT-ELISA device. Prior to fabricating the full device, we first optimized the individual mixing, depletion, and detection modules. The mixing module achieves rapid and continuous mixing of reagents through the use of serpentine channels with optimized length and incorporates herringbone structures inside the channel^31,32^ (Figure 3A, bottom). This dramatically enhances the rate of molecular diffusion and reduces the required incubation time to less than one minute. We injected different concentrations of insulin into the mixing module through the sample inlet and injected a solution of fluorescently tagged detection antibodies and microbeads functionalized with the insulin capture antibody through the reagent inlet. We injected these solutions at different flow rates, and then collected the beads from the module outlet and observed them under a fluorescence microscope. After comparing the fluorescence signal intensity obtained at different flow rates with the intensity achieved with standard bench-top incubation (Figure 3A, top), we found that optimal mixing occurs at a flow rate of 15 µL/min, with a total mixing time of 30 sec. At lower or higher flow rates, mixing is not as effective as bench-top incubation.

**Figure 3.**
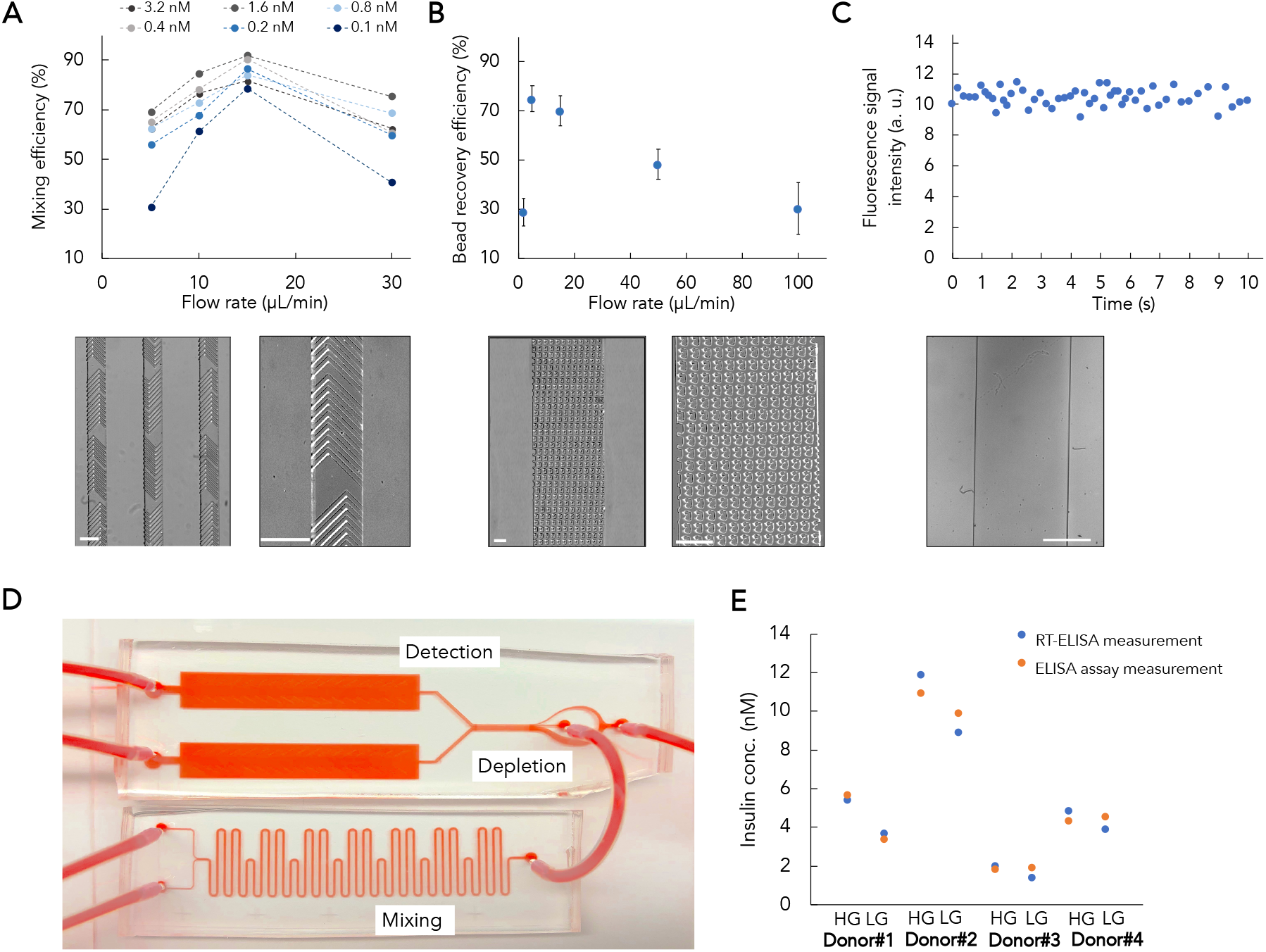
Testing and optimizing the core components of RT-ELISA. **A)** Optimizing the performance of the mixing module. We injected different concentrations of insulin and detection reagents through their respective inlets at different flow rates. We then collected the beads and observed their fluorescence signal under a microscope relative to a detection reaction with standard two hours bench-top incubation (top). An image of the mixer device with serpentine channel and herringbone structures (bottom). **B)** Optimizing the performance of the depletion module. We spiked whole blood with 15 μm microbeads and tested the performance of notched post structures for bead recovery and blood cell depletion at different flowrates (top). An image of the depletion module is shown at bottom. **C)** Optimizing the performance of the detection module and assessing the temporal resolution. We introduced fluorescently labeled microbeads into the detection module and observed the beads over 10 s. We measured 5 beads per second on average passing through the detection window, which defines a temporal resolution of 200 ms (top). An image of the detection window is shown at bottom. **D)** The fully integrated RT-ELISA device. **E)** Insulin probes were incorporated into the device and used to measure endogenous insulin secreted from human islet samples after a glucose challenge: high glucose (HG) or low glucose (LG). The RT-ELISA readout was compared to conventional ELISA. The experiment was performed three times with human islets from four donors. White and black scale bars show 150 µm and 5 mm, respectively.

The depletion module is designed to not only eliminate blood cells, but also free fluorescently tagged antibodies, thereby reducing the background (**Video S1**). This module (Figure 3B) employs deterministic lateral displacement (DLD) sorting to isolate beads from blood cells. DLD is an established hydrodynamic approach for separating particles based purely on size in a continuous manner^33^. DLD utilizes specific arrangements of posts within a channel to facilitate separation of particles larger and smaller than a critical diameter (D_c_) by precisely controlling their trajectory within the device. In this scenario, the D_c_ is 15 µm, based on the diameter of the microbeads used for target capture, and we sought to design a device that would allow us to isolate these beads amid a far larger number of blood cells, which are smaller than 15 µm. We tested notched^34^ post structures (see Methods for details on depletion module design), and introduced a sheath buffer solution into the device along with the sample for bead recovery and blood cell depletion. We achieved the best performance (both for bead recovery and blood cell depletion) at a sample flow rate of 5-20 µL/min and a sheath buffer to sample flow-rate ratio of 5:1 (Figure 3B, top; **Figure S1-2**). At higher or lower inlet flow rates, the depletion performance worsened.

In the detection module, the target-bound beads flow into a detection window (Figure 3C, bottom), and their fluorescence intensity is continuously measured with a high-speed camera under spatially multiplexed two-color laser illumination (see Methods and Supplementary Information). Incoming beads are illuminated by a red laser first, which interrogates the Cy5 fluorescence intensity that indicates glucose concentration. This is followed by illumination with a green laser that interrogates the R-PE fluorescence intensity, which measures insulin concentrations. We used an exposure time of 50 ms and acquired images every 100 ms (**Video S2**). We introduced fluorescently labeled microbeads into the detection module at the inlet flow rate of 15 µL/min and measured the fluorescence signal intensity and the number of beads passing through the detection window over a short time scale. On average, we observed 5 beads per second passing through the detection window, indicating the temporal resolution of approximately 200 ms (Figure 3C, top). The fast framerate and high sensitivity of the camera allow us to continuously track individual microbeads, enabling quantitative detection of analytes in real time. Finally, we combined the three modules to produce the integrated RT-ELISA device (Figure 3D). To achieve this, we simulated and adjusted the fluidic resistance between the different modules. The process used to generate the fully integrated device is described in the Methods and **Figures S3-5**.

Next, we tested the RT-ELISA device for detecting endogenous, secreted human insulin^35^. We treated intact human islet with culture media containing low (2.8 mM) and then high (16.7 mM) concentrations of glucose to stimulate insulin secretion *in vitro*. After each stimulation, the supernatants were collected, and insulin concentration was measured using both the RT-ELISA device and conventional ELISA. We performed this experiment using islets from four different donors. The results presented in Figure 3E shows that our bead-based assay is capable of measuring endogenous human insulin levels, and that the measurements obtained from our device closely matched those from the ELISA assay (**Figure S9**).

### Continuous measurement of glucose and insulin in whole blood

To characterize the sensitivity and temporal resolution of the RT-ELISA device, we tested it with human whole blood spiked with known concentrations of glucose and insulin (Figure 4). We used these data to construct standard curves that correlate fluorescence signal intensity to glucose or insulin concentration in whole blood (Figure 4A, B, insets).

**Figure 4.**
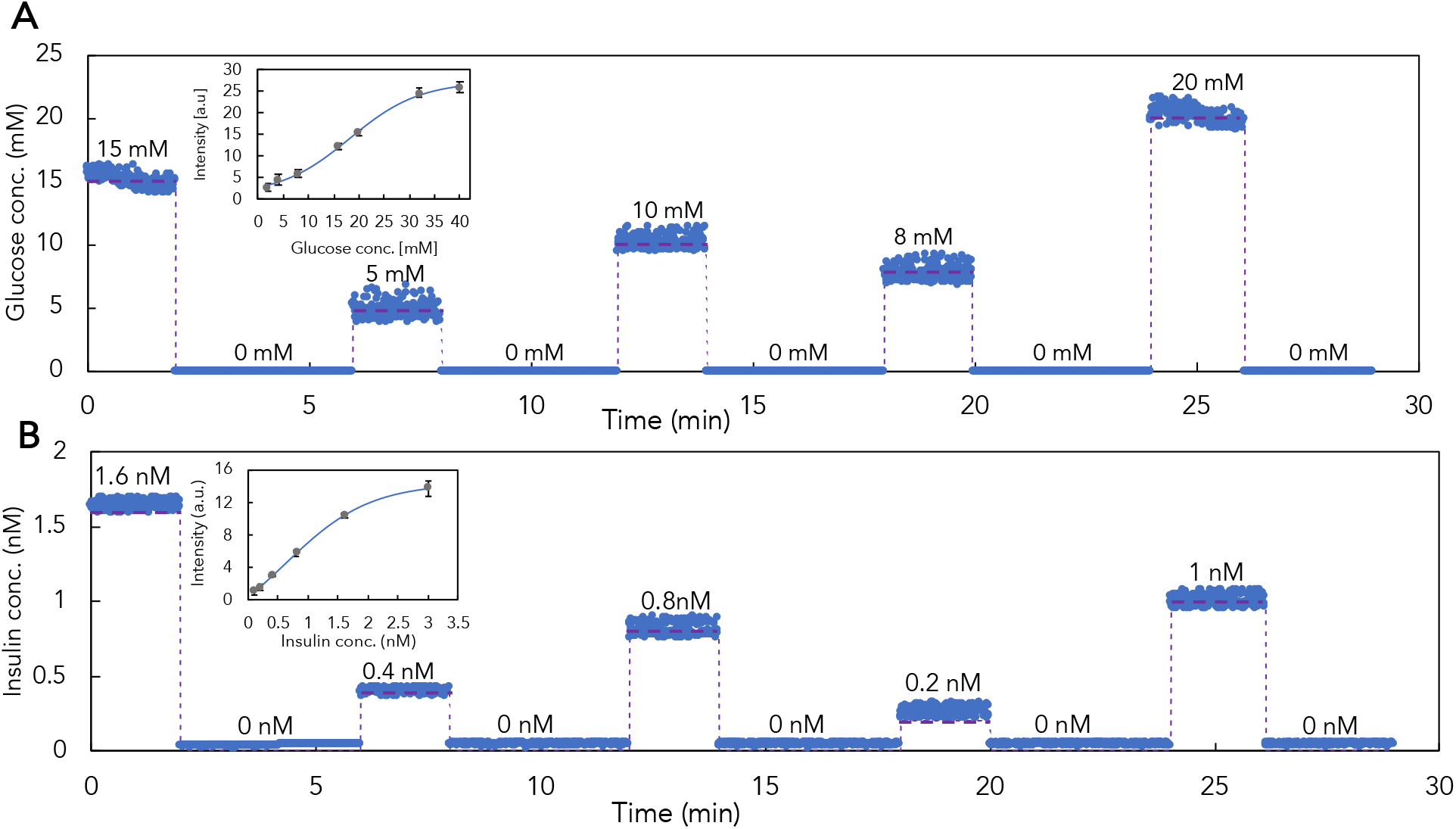
Continuous *in vitro* monitoring of glucose and insulin in whole blood. RT-ELISA measurements of glucose (**A**) and insulin (**B)** in human whole blood (blue dots) relative to actual concentrations (purple dashed line) over the course of 30 mins. Insets show standard curves correlating fluorescence signal intensity to glucose or insulin concentrations. Experiments to derive standard curves were repeated three times (replicates shown in **Figures S6-7**), and error bars show the standard deviation. Non-linear regression analysis is used for curve-fitting. Data reflect individual bead readouts, where at least 200 beads were measured for each concentration. We used the following concentrations in these experiments for glucose: 15 mM, 5 mM, 10 mM, 8mM, and 20 mM; and for insulin:1.6 nM, 0.4 nM, 0.8 nM, 0.2 nM, and 1 nM. Measurements were performed for 2 mins; between sample changes, buffered solution was injected into the device for 4 mins. The fluorescence signal intensity from a blank blood sample was measured to quantify endogenous insulin and glucose. For insulin measurements, the fluorescence signal intensity from the blank was subtracted from the measured signal at each concentration. It is worth mentioning the endogenous insulin concentration in the blood was significantly lower than all concentrations measured during subsequent experiments and therefore we neglected the endogenous component in making the calibration curve. For glucose measurements, endogenous glucose was measured using a conventional glucose meter and subtracted as baseline before calculating the spiked concentrations.

We developed custom software to continuously measure the fluorescence signal intensity and quantify glucose and insulin concentrations (see Methods). Our device achieved a LOD of 3.7 mM for glucose and 93 pM for insulin measurements in blood. Although fasting insulin in non-diabetic patients (~40 pM) is below this limit, an LOD of 93 pM is sufficient to track the pharmacokinetics of post-prandial insulin. Although the LOD will need to be further optimized to detect hypoglycemia (LOD=2.2 mM)^36^, our glucose detection limits cover both the euglycemic range and relevant hyperglycemic range. We then introduced different concentrations of glucose and insulin in a whole blood sample and used the RT-ELISA device to monitor these analytes for approximately 30 mins (Figure 4). The measured concentrations differed from the known concentration on average by 250 μM and 24 pM for glucose and insulin measurements, respectively, over the course of the experiment. This standard deviation is sufficiently small that our system can be employed to reliably quantify glucose and insulin in diabetic patients, where post-prandial concentrations are typically more than 3.5 mM and 300 pM, respectively.

### Continuous glucose and insulin measurements *in vivo*

Having demonstrated our platform’s ability to sensitively and accurately detect insulin and glucose *in vitro*, we evaluated its performance *in vivo* in a streptozotocin-induced rat model of insulin-deficient diabetes^37^. We chose this model because it lacks endogenous insulin, eliminating the potential of background due to antibody cross-reactivity. The device was connected to an anesthetized rat by a femoral venous catheter (**Figure S11**). Humulin R (recombinant human insulin) boluses were given at one (t=0) or two timepoints (t=0 and 20 min), while continuous, real-time monitoring of glucose (Figure 5A, C, E) and insulin (Figure 5B, D, F) levels were performed over the course of 30 to 50 mins. In experiments monitoring only a single insulin bolus (Figure 5A, C), baseline insulin measurements were elevated due to insulin boluses given 40 minutes prior to the experiment. Peak insulin concentrations were detected at either 5 min (Figure 5B) or 10 min (Figure 5D) after single insulin injections. After two successive bolus injections, peak insulin concentrations were observed at 10 and 35 min (Figure 5F). In parallel, we compared the RT-ELISA results with conventional ELISA analysis of insulin in blood samples collected from the rat tail vein every 5 min and glucose measurements from a handheld glucose monitor and observed that both sets of results correlate well (**Figure S10**). The differences between peak insulin action in individual rats clearly show the variability in insulin pharmacokinetics, even under controlled conditions with genetically similar animals. This has important implications for human patients, who are genetically diverse, use different injection sites, and have different environmental conditions, exacerbating this inter-individual variability. Thus, these results highlight the necessity of personalized insulin monitoring.

**Figure 5.**
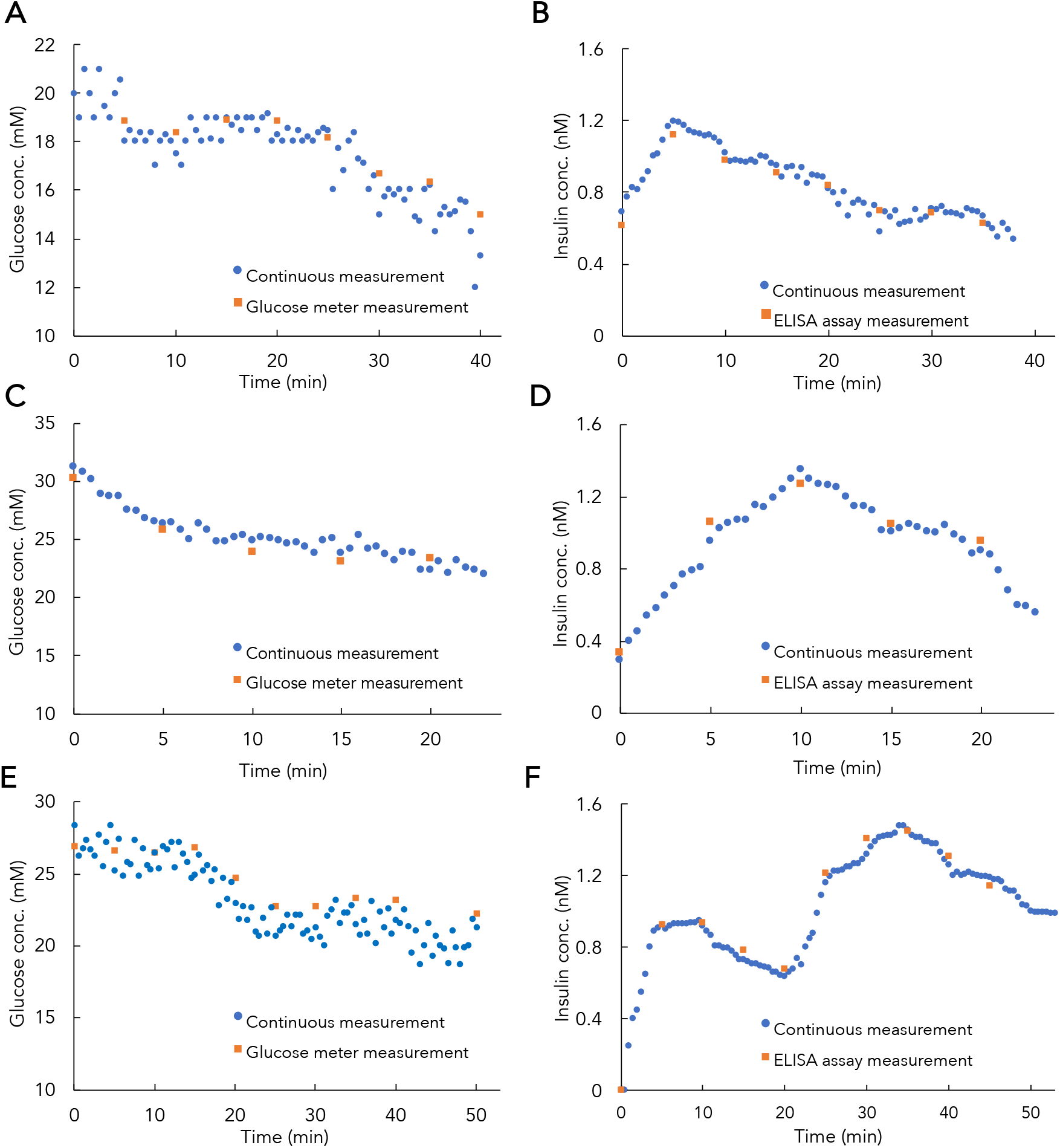
Continuous, real-time measurements of glucose and insulin in diabetic rats. RT-ELISA measurement of *in vivo* glucose **(A, C, E)** and insulin **(B, D, F)** concentrations over 30-50 mins in three different rats. Diabetic rats were injected subcutaneously with a single bolus of Humulin R (1U/kg) at t = 0 (**A**-**D**) or two boluses at t=0 and 20 min (**E, F**). Each blue dot shows the median RT-ELISA readout of individual bead (~150 beads) measurements over 30 sec. For comparison, we collected blood samples from the tail vein every 5 min, and measured glucose and insulin levels via handheld glucose meter and conventional ELISA, respectively (orange squares). These results correlated closely and highlight the inter-individual variability in insulin response.

We observed that the insulin dose used in our experiments is not sufficient to decrease blood glucose concentrations to normoglycemia conditions. However, we demonstrate that the observed decrease in blood glucose after insulin administration in our anesthetized rats is comparable to blood glucose depletion in a conscious rat. We administered the same insulin dose (1U/kg) to a conscious rat and collected blood samples at t = 0, 5, 20, and 40 min after injection. We then employed the RT-ELISA device to measure glucose (Figure 6A) and insulin (Figure 6B) levels. We observed a decrease in glucose concentration both in the glucose meter measurements and in the RT-ELISA device, with a 7 mM decrease in glucose level after 40 minutes. These results are comparable to the results from the continuous monitoring experiments and corroborate the expected effect of a 1U/kg insulin bolus on glucose concentrations.

**Figure 6.**
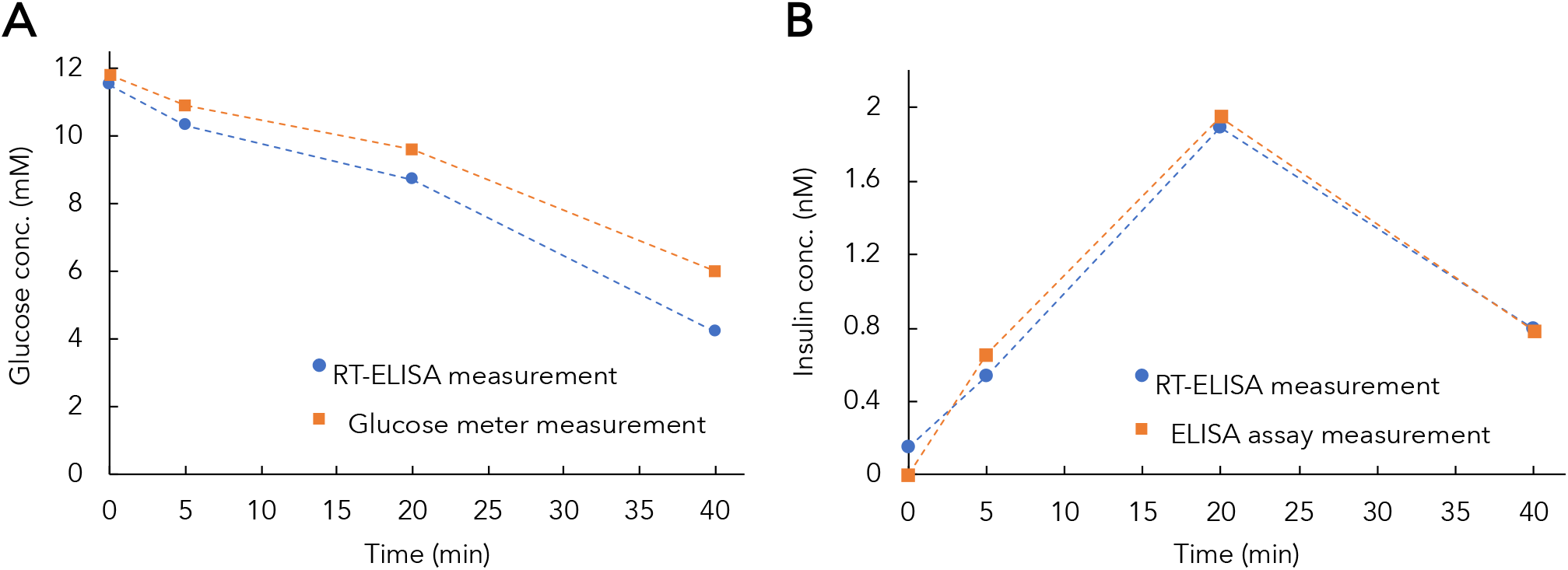
Glucose and insulin measurements in an awake diabetic rat. An awake diabetic rat was injected with human insulin, with blood samples collected at t = 0, 5, 20, and 40 min after injection. Glucose levels were measured using RT-ELISA and glucose meter **(A)**, and insulin levels were measured using RT-ELISA and ELISA assay **(B)**.

### Continuous *in vivo* tracking of different insulin formulations

There are both short and long-acting commercially available recombinant insulin formulations, which exhibit different absorption kinetics after injection. We have found that RT-ELISA can differentiate the pharmacokinetics between these formulations, based on experiments comparing the kinetics of Humulin R and Humulin N (neutral protamine Hagedorn insulin). Humulin R is considered a short-acting insulin, while Humulin N was developed to achieve delayed onset and extended duration of action as an early basal insulin replacement^41–43^.

After measuring baseline concentrations for 5 mins, diabetic rats were injected with Humulin R or Humulin N and glucose and insulin concentrations were quantified over the course of 85 mins (Figure 7 **and Figure S8**). The different pharmacokinetics of Humulin R and Humulin N were clearly evident in the RT-ELISA measurements, with these two formulations producing peaks at approximately 15 mins and 60 mins after injection, respectively.

**Figure 7.**
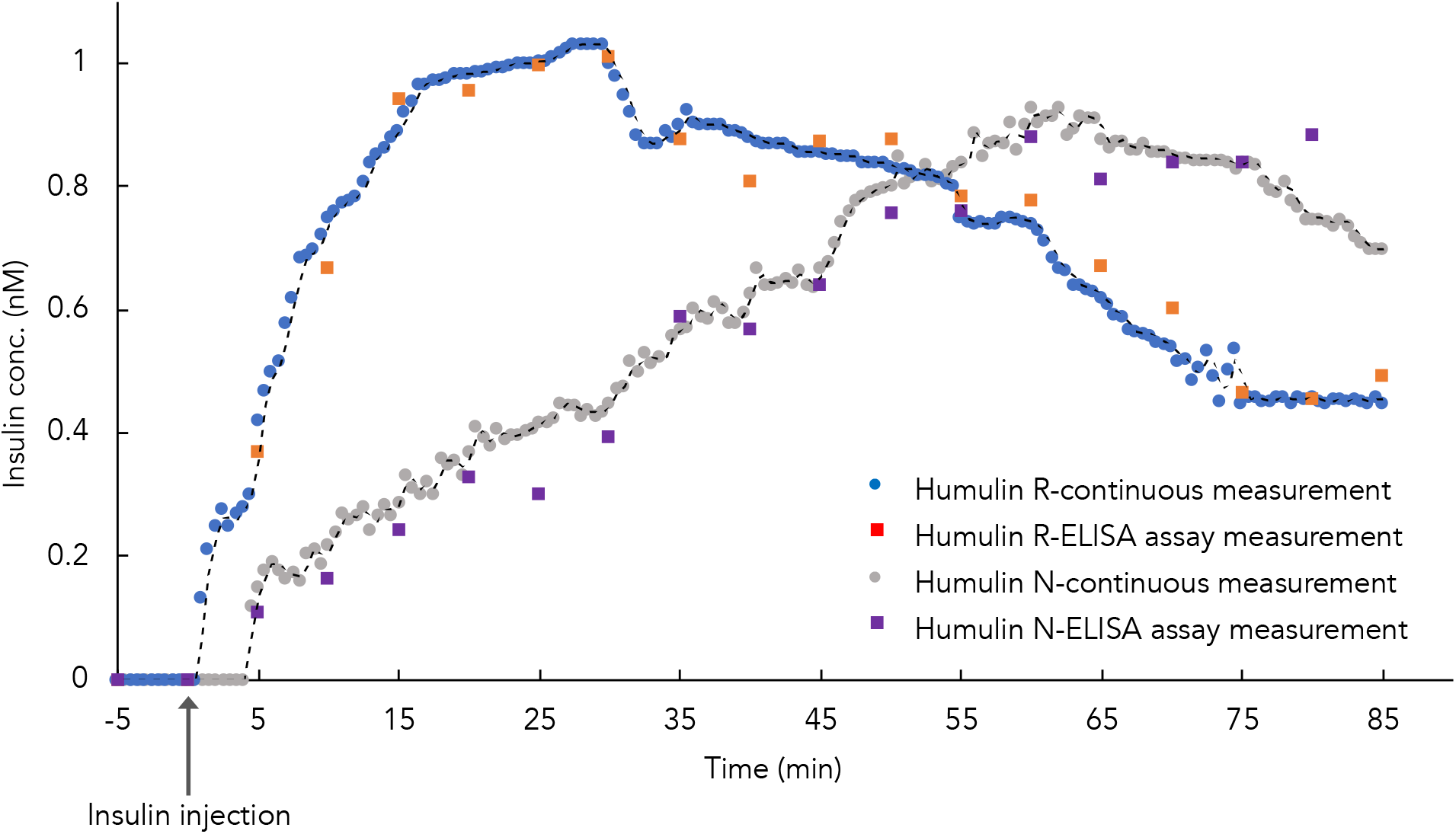
Comparison of different insulin formulation pharmacokinetics. Diabetic rats were injected with either Humulin R (1U/kg) or Humulin N (2U/kg), and insulin concentrations were monitored using RT-ELISA and ELISA.

## Conclusion

We have demonstrated that the RT-ELISA device can achieve sensitive and accurate continuous monitoring of insulin and glucose in the circulating blood of live animals, enabling real-time *in vivo* analysis of insulin pharmacokinetics and pharmacodynamics. Experiments in human blood demonstrated that our sensor achieved sufficient sensitivity to detect clinically relevant concentrations of both analytes, with a LOD of 3.7 mM for glucose and 93 pM for insulin. Subsequent experiments in anesthetized diabetic rats confirmed our capacity to continuously monitor *in vivo* changes in insulin and glucose in real-time and highlighted the RT-ELISA platform’s capacity to distinguish clear inter-individual differences in the pharmacokinetics of insulin between animals—a critical feature for clinical implementation. Importantly, RT-ELISA measurements closely matched those obtained with standard ELISA and clinical glucose sensors; we noted with both readouts that the use of isoflurane anesthesia impeded the response of rats to insulin, but subsequently confirmed our ability to accurately track insulin-induced changes in circulating glucose with samples collected from awake rats at multiple timepoints. Finally, we demonstrated the capacity to accurately discriminate the distinct pharmacokinetic profiles associated with two different insulin formulations—short-acting Humulin R, and intermediate-acting Humulin N.

Although it is beyond the scope of this work, we believe our system could be readily modified for human use. Important considerations to that end would include careful preparation of tubing and the chip surface with anticoagulants to avoid clotting as well as safety measures to ensure that reagent solutions cannot flow back into the bloodstream. With such modifications, RT-ELISA could provide the foundation for a clinically useful assay for developing personalized insulin therapy regimens for individual patients, and it could also serve as a valuable research tool for assessing the pharmacokinetics and pharmacodynamics of novel insulin formulations for both preclinical testing and clinical trials.

While in this work we demonstrate the simultaneous detection of two analytes, we believe that higher levels of multiplexing can be achieved with a more advanced optical design. Finally, we emphasize that the RT-ELISA system is a platform technology that could be readily modified to continuously measure other circulating analytes *in vivo*, for which aptamers or antibody pairs are available, thus making it potentially a versatile tool for biomedical research.

## Materials & Methods

### Materials

All chemicals were purchased from Sigma-Aldrich, unless otherwise noted. 15-µm SuperAvidin-coated microspheres were obtained from Bangs Laboratories, human whole blood was obtained from BioIVT, matched paired insulin antibodies (capture: BM364-Z8A2, detection: BM364-T8F5) were obtained from BBI Solutions, human insulin was obtained from R&D Systems, Streptozotocin (STZ) was obtained from MedChem Express, Humulin R was obtained from Eli Lilly, Humulin H was obtained from Eli Lilly, R-PE antibody conjugation kit was obtained from TriLink Biotechnologies, and the glucose aptamer and DNA competitor oligonucleotides (see Table 1 for sequences) were ordered from Integrated DNA Technologies.

**Table 1.**
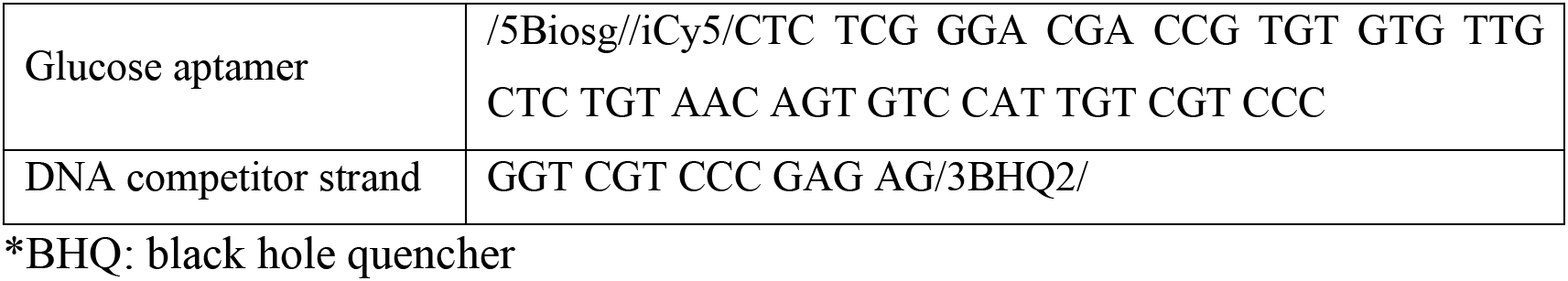
Sequence of the nucleic acids, utilized for glucose monitoring

### RT-ELISA device fabrication

A polydimethoxysilane (PDMS; Dow Chemical) device fabrication protocol was employed. First, two silicon masters were fabricated: one for the mixing module and one for the depletion and detection modules. Masks were designed using AutoCAD software (**Figure S5**) and then printed (CAD/Art Services & Advance Production Inc.). Two 50-μm layers of SU8-3050 (Microchem) were patterned to make the mixing module master. The first layer was patterned to make the channel, and the second was aligned for herringbone structure fabrication. The depletion and detection module masters were patterned using one layer of SU-8. After master fabrication, PDMS and curing agent with the ratio of 10:1 were poured onto the device masters. Both masters were baked at 67 °C for 2.5 hours. Afterward, we peeled the replicas and pierced holes to connect the tubing. PDMS replicas were attached to glass cover slips using a 45 sec plasma treatment and left to bond overnight. The silicon tubing was then attached to the inlet and outlet of the device. Prior to use, devices were conditioned with 1% Pluoronic F68 (Sigma-Aldrich) and 10% heparin (McKesson Corporation) in phosphate-buffered saline (PBS) overnight in order to reduce nonspecific adsorption and blood coagulation.

### Depletion module design

We tested notched pillar structures in the depletion module. Pillar array positioning was calculated for the separation of particles larger and smaller than 14 μm. This cutoff was chosen to ensure the maximum recovery of 15-μm microbeads while depleting the majority of blood cells, based on the critical diameter defined by the empirical formula for DLD ^33^. These parameters were then relaxed by adjusting gaps according to notched pillar shapes. The DLD arrays extended across the entire width of the device and were then trimmed in AutoCAD to create continuous walls. The sheath buffer inlet splits so that it enters along both edges of the device, ensuring that no particles or cells enter the device next to a wall. Different sample inlet to buffer inlet flow-rate ratios were examined, with optimal cell recovery and blood cell depletion observed at a ratio of 1:5. **Figure S1** shows the separation of microbeads from blood cells. Prior to separation, blood cell background blocks signal from the beads, whereas beads can be clearly detected after separation. The optimal pillar diameter to maximize sorting efficiency in a DLD device with notched microstructures was 40 µm, based on calculations described in the literature^33^.

### Integrated RT-ELISA device fabrication

We first connected the mixer and depletion modules and tested their performance in capturing insulin from whole blood samples and isolating target-bound beads. As shown in **Figure S3**, blood spiked with 3 nM insulin was injected through the sample inlet at 15 μL/min, while the reagent solution containing functionalized microbeads and detection antibodies was injected through the reagent inlet at the same flowrate. The mixer outlet was connected to the depletion sample inlet, and as the mixture entered the depletion module, a buffered solution was injected at 75 μL/min to form the required sheath flow for separation. The flowrates were optimized to achieve the best mixing efficiency, bead recovery, and blood cell depletion. The beads were collected from the depletion outlet and monitored via fluorescence microscopy.

The connection of the depletion and detection modules was tested prior to full integration. A dummy detection module was connected to the blood waste outlet to adjust outlet resistance, ensuring fluid flow through the depletion device (**Figure S4**). The detection window was designed to have a width of 292 μm in order to accommodate the 20X objective field used for fluorescence measurement. The detection module was then widened to 4.8 mm, to prevent clogging by the high number of blood cells and beads passing through the channel. The optimized dimensions of the modules in the final device are summarized in **Table S1**.

### Microbead functionalization

Streptavidin-coated beads were functionalized with either monoclonal biotinylated anti-insulin capture antibodies (cAbs) or glucose aptamers hybridized with the DNA competitor strand, as directed by the bead manufacturer’s recommended protocol. A typical coating procedure for 10 mg of microspheres entailed homogenizing the bead stock solution on a rotator, transferring 1 mL of the 10 mg/mL stock to a new tube, adding 10 mL of 1x PBS + 0.05% Tween 20 (PBST, pH 7.4), mixing by pipetting or vortexing, centrifuging at 2100 rcf for 5 min, and removing the supernatant. The second wash was performed in the same manner. Beads were then resuspended in 20 mL PBS + 0.05% BSA + 0.01% Tween 20 (PBSBT) for insulin beads and PBSBT + 100 mM MgCl_2_ (PBSBTMg) for glucose beads, and incubated with 1.2 nM biotinylated cAb, such that the cAb was present in excess (1.5-2 µg cAb/mg beads) compared to the amount required to form a monolayer on the surface of each bead. The mixture was briefly vortexed or pipetted to homogeneity and incubated at room temperature on a rotator for 30 min. Following this, the beads were pelleted via centrifugation at 2100 rcf for 5 min and washed thrice with 10 mL PBST as described above. The supernatant was removed, and the beads were resuspended in PBSBT or PBSBTMg at a concentration of 10 mg/mL and stored at 4°C until used.

### Insulin fluorescence-based sandwich immunoassay

The detection antibody (dAb) was site-specifically labeled with R-Phycoerythrin (R-PE) using a SiteClick R-PE Antibody Labeling Kit (Thermo Fisher Scientific) according to the kit manufacturer’s instructions. We employed the dAb conjugated with R-PE along with the bead-coupled cAbs in either bench-top incubation or the RT-ELISA device to achieve insulin detection.

### Imaging setup

We used a compact home-built fluorescent microscope with spatially multiplexed two-color laser illumination and wide-field high-speed fluorescent imaging to measure the fluorescence of the beads. A Nikon CFI Plan Apo VC 20X Air 0.75 NA UV objective was used to focus two diode lasers (520 nm/40 mW and 642 nm/5 mW) with proper excitation filters (Thorlabs), to a size of ~500 by 100 µm at the sample plane. The two laser spots have a spatial offset of ~500 µm along the bead flow direction to enable spatially multiplexed two-color fluorescent readout. The entire field of view covering both spots was imaged continuously by a high-speed, high-sensitivity scientific sCMOS camera (Photometrics Prime 95B) after proper two-pass-band emission filters (Chroma). Incoming beads are illuminated by the 642-nm laser first, which interrogates the Cy5 fluorescence intensity that indicates glucose detection, followed by the 520-nm laser, which interrogates the R-PE fluorescence intensity that indicates insulin concentrations. To ensure that red (green) laser does not excite R-PE (Cy5) fluorophore, we performed control experiments where only insulin (glucose) sample and its reagent solution were injected while both lasers were illuminating. We observed insulin (glucose) beads passing through the green (red) laser region while no signal was observed in red (green) laser region. A schematic of the imaging setup is provided in **Figure S12**.

### *In vitro* glucose-stimulated insulin secretion assays

Deidentified human pancreatic islets were obtained from four previously healthy, non-diabetic organ donors through the Integrated Islet Distribution Program (http://iidp.coh.org), Alberta Diabetes Institute Islet Core and University of California San Francisco Islet Core. 150 primary human islets were used for each experiment. Secretion assays were carried out at 37°C in RPMI 1640 (Thermo Fisher Scientific) media supplemented with 0.5% (v/v) fetal bovine serum (HyClone), 0.2% (w/v) bovine serum albumin (Sigma-Aldrich) and D-glucose (Sigma-Aldrich) at indicated concentrations. In brief, islets were incubated at a glucose concentration of 2.8 mM for two sequential incubations, each for 45 minutes, (media is changed after each incubation) as initial equilibration period. Subsequently, islets were challenged with 2.8 mM (low) and then 16.7 mM (high) glucose at 37 °C for 60 min. Supernatants were carefully collected at the end of challenge for insulin quantitation by either a conventional human insulin ELISA kit (Mercodia) or RT-ELISA.

### Derivation of the LOD

In order to estimate the LOD of the RT-ELISA device for glucose and insulin measurements, we first measured the fluorescence signal intensity from the blank sample and calculated the *FL*_*L*_:

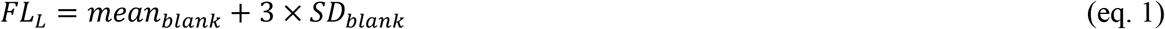

We then generated a standard curve to calculate the LOD^44^.

### Continuous, *in vitro* monitoring of glucose and insulin in whole blood

Human blood samples were spiked with different concentrations of glucose and insulin for *in vitro* tracking. Blood glucose was measured using a handheld glucose meter (Bayer Contour Next) to calculate the spiked glucose concentrations. A reagent solution consisting of cAb-functionalized beads, PE-tagged dAbs, and glucose aptamer-capture strand complex-functionalized microbeads was prepared in buffered solution (1x PBS + 0.05% BSA + 0.01% Tween 20 + 100 mM MgCl_2_). Different concentrations of glucose and insulin, respectively—15mM + 1.6 nM; 5 mM + 0.4 nM; 10 mM + 0.8 nM; 8 mM + 0.2 nM; or 20 mM + 1 nM—were spiked into whole blood and injected into the device sequentially. Spiked blood samples and the reagent solution were injected through the inlets of the mixer device at a flowrate of 15 μL/min. Between changing samples, buffered solution was injected into the device for 4 mins. A blood sample with no spiked insulin or glucose was run through the device to measure the fluorescence signal intensity from endogenous insulin, and this was subtracted from the measured signal at each concentration. At the end of experiment, the images were collected, and a custom-written program was used to measure the fluorescence signal intensity, as described in the following section. The derived standard curves were used to calculate absolute concentrations.

### Image analysis

As a proxy to measure the intensity of the glucose and insulin molecules, we measured the intensity of the signal from beads passing through the detection window, which was carried out in three steps. First, we measured the background intensity. This was done by constructing an intensity profile image containing the median intensity of each pixel across all recorded frames at different time stamps. We then applied a smoothing Gaussian filter with a small 5 x 5 pixel kernel to this image to produce the background intensity. Second, we localized the beads. We subtracted the background intensity from each frame, and then sought for connected high-intensity components within each frame. We used a (tunable) threshold on the intensity level to obtain a binary mask and applied a second (tunable) threshold on the size of the connected component to localize the beads in each frame. Finally, we computed the intensity profile over all beads, and computed statistics such as mean, median, and standard deviation. We exported this data into a CSV file for further processing (*e.g.*, using Excel). We also produced a video localizing the beads. We determined the edges of each bead from step 2 and processed the frame to highlight the edges of the localized beads. We saved all of these processed frames in a video (avi format). This provides a way to visualize the detected beads, and also could be used as a way to tune the bead localization thresholds in step 2. These steps were implemented in Python, and the code and Jupyter notebook for these steps and performing the images analysis will be released on GitHub.

### STZ-induced model of diabetes in rats

Animal studies were performed in accordance with the Guidelines for the Care and Use of Laboratory Animals and the Animal Welfare Act Regulations; all protocols were approved by the Stanford Institutional Animal Care and Use Committee. The protocol used for STZ induction was adapted from the protocol by Kenneth K. Wu and Youming Huan^37^. Briefly, male Sprague Dawley rats (Charles River Laboratory) of 160–230 g (8-10 weeks) were weighed and fasted 6-8 hours prior to treatment with STZ. STZ was diluted to 10 mg/mL in sodium citrate buffer immediately before injection. STZ solution was injected intraperitoneally at 65 mg/kg into each rat. Rats were provided with water containing 10% sucrose for 24 hours after injection with STZ. Rat blood glucose levels were tested for hyperglycemia daily after the STZ treatment via tail vein blood collection using a handheld Bayer Contour Next glucose monitor. Diabetes was defined as three consecutive blood glucose measurements >400 mg/dL in non-fasted rats.

### Continuous monitoring of glucose and insulin in diabetic rats

Diabetic rats were fasted for 4-6 hours. Rats were anesthetized using 1–3% isoflurane and a 20G catheter was inserted into the femoral vein. The catheter was then connected to a peristaltic pump (Ismatec) at a flow rate of 15 μL/min. Once the tubing was primed with blood, the device was connected, and the rats were injected subcutaneously with either 1 U/kg Humulin R or 2 U/kg Humulin N. As the blood entered the device, reagent solution was injected through the reagent inlet. Images were collected from the detection module and the fluorescence signal intensity was quantified as described above, and glucose and insulin absolute concentrations were calculated. After injection, blood was sampled every 5 mins for up to 120 mins. Before injection, baseline blood glucose was measured using a handheld glucose meter, and blood was collected in serum tubes (Starstedt) for analysis with a commercial human ELISA kit (Mercodia).

## Acknowledgements

This research was supported by the Chan-Zuckerberg Biohub and the Stanford Diabetes Research Center (SDRC) Pilot grant. C.L.M. was supported by the NSERC Postgraduate Scholarship and the Stanford Bio-X Bowes Graduate Student Fellowship. We are thankful to Bruce Buckingham and Rayhan Lal for their thoughts and helpful discussions. We also thank Nicolo Maganzini and Ian Thompson for their review and edits on the manuscript. We acknowledge Stanford Nanofabrication Facility (NSF) for their cleanroom facilities. The authors thank the Stanford Veterinary Service Centre staff for their assistance with animal care and procedures.

## Author contributions

M.P., C.L.M., and H.T.S. conceived the initial concept. M.P. and C.L.M. designed experiments. M.P., C.L.M., E.Y.M., J.P., D.M., Y.H., and S.W.B. executed experiments. A.B. developed the imaging analysis algorithm. M.P. and C.L.M. analyzed the data. M.P., C.L.M., M.E., and H.T.S. wrote the manuscript. All authors edited, discussed, and approved the whole paper.

## Additional information

Online Supplementary Information accompanies this paper.

