## Supplementary Information for "Continuous detection of glucose and insulin in live animals"

**This PDF file includes:**

**Table S1.** Dimensions of fabricated modules of the RT-ELISA device

**Figure S1.** DLD characterization
**Figure S2.** Simulated flow velocity inside a depletion module with a notched pillar structure
**Figure S3.** Integration of the mixing and depletion modules
**Figure S4.** Integration of the depletion and detection modules
**Figure S5.** Integrated RT-ELISA platform mask design
**Figure S6.** Replicates of glucose standard curve derivation
**Figure S7.** Replicates of insulin standard curve derivation
**Figure S8.** Glucose monitoring data in live rats after injection of two different insulin formulations.
**Figure S9.** Regression statistical comparison of RT-ELISA and ELISA for *in vitro* secretion assays
**Figure S10.** Regression statistical comparison of RT-ELISA and ELISA or glucose monitor for *in vivo* rat measurements.
**Figure S11.** Animal experiment setup
**Figure S12.** Schematic of imaging setup

**Microscope videos**

**Table S1. Dimensions of fabricated modules of the RT-ELISA device**

| | |
|---|---|
| **Mixer module** | |
| Sample & reagent inlet width | 150 μm |
| Channel width | 300 μm |
| Outlet width | 150 μm |
| Serpentine channel length | 30 cm |
| **Depletion module** | |
| Sample inlet width | 150 μm |
| Total channel width | 850 μm |
| Channel length | 7.5 mm |
| Pillar to pillar distance (λ) | 55 μm |
| Gap (horizontal) | 30 μm |
| Gap (vertical) | 22 μm |
| Periodicity [# rows] | 10 |
| Offset between pillar rows | 11 degrees |
| Theoretical critical diameter | 14 μm |
| **Detection module** | |
| Small (main) channel width | 292 μm |
| Large channel width | 4800 μm |

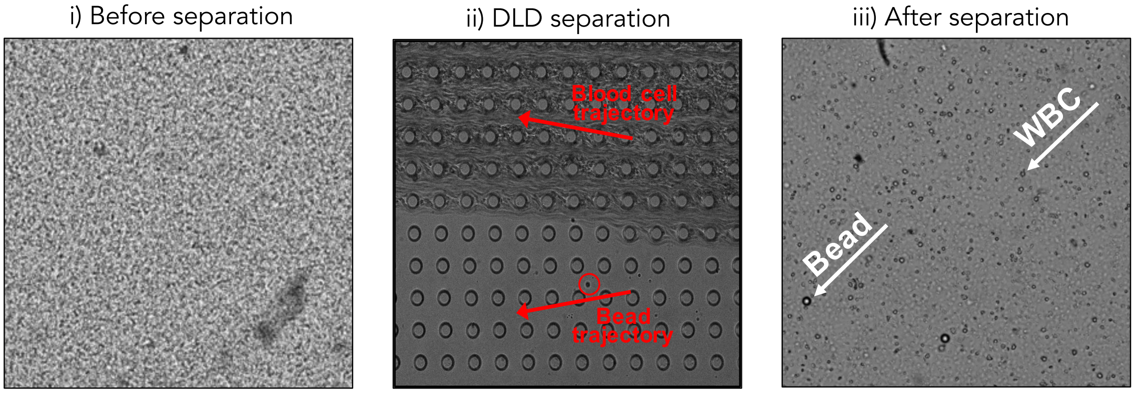


**Figure S1. DLD characterization.** Images of a blood sample spiked with 15-µm beads before, during, and after DLD separation using circular pillar structures.

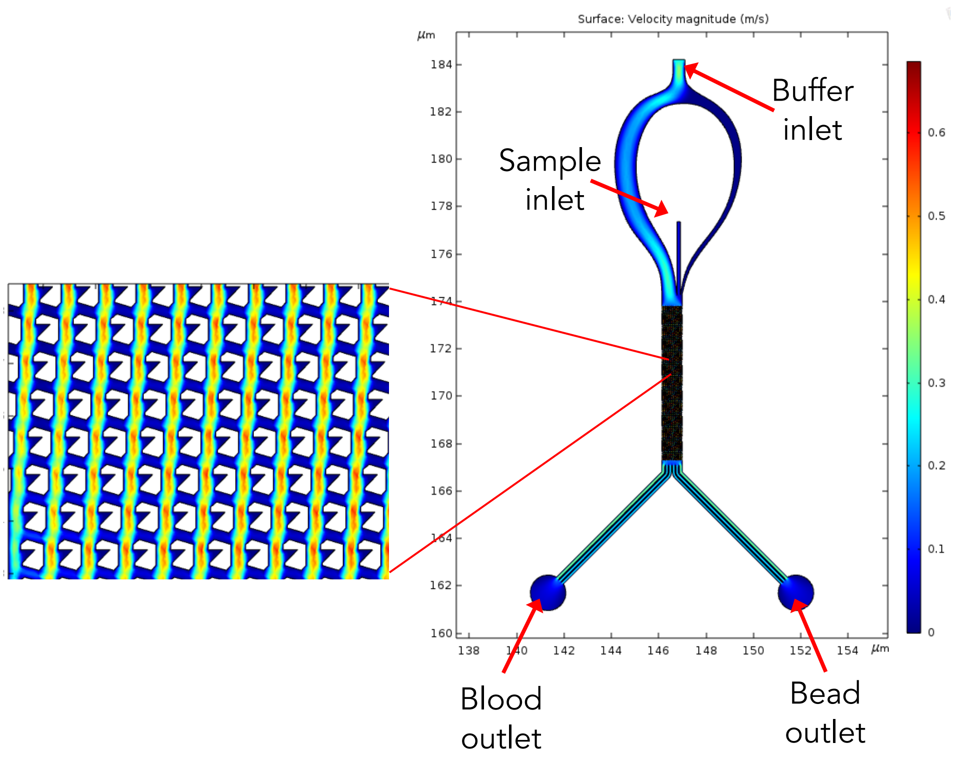


**Figure S2. Simulated flow velocity inside a depletion module with a notched pillar structure.** Flow velocity was simulated inside the depletion module using COMSOL Multiphysics software.

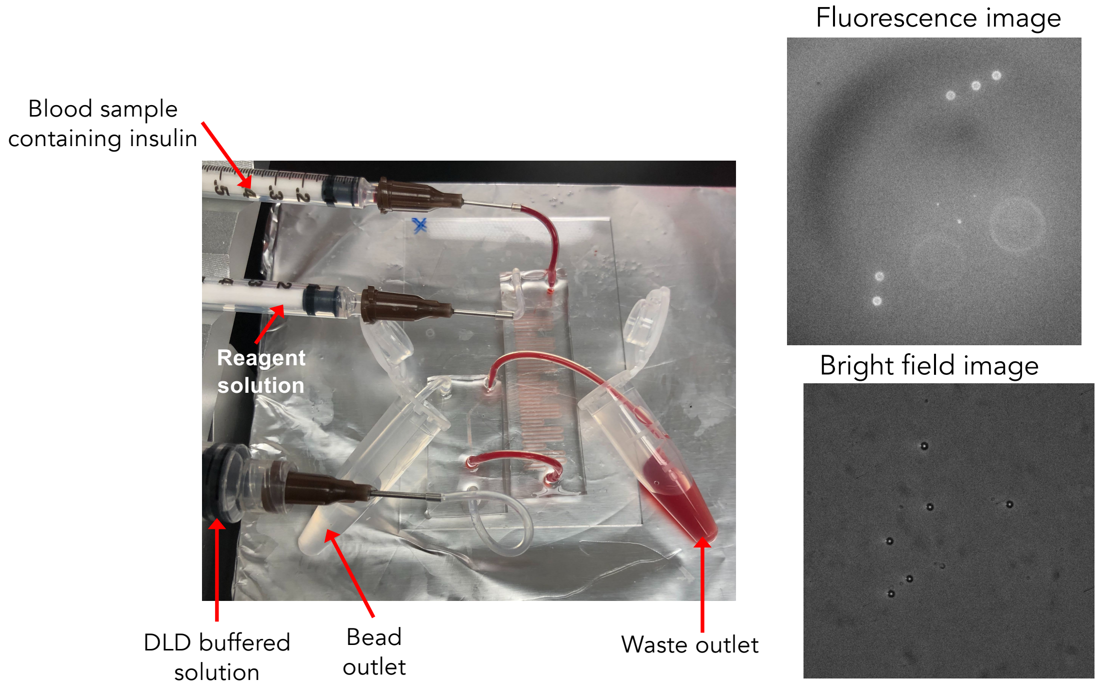


**Figure S3. Integration of the mixing and depletion modules.** The mixing module was connected to the depletion module through tubing. A whole blood sample spiked with 3 nM insulin and the detection reagent solution were injected through their respective inlets. The beads were collected from the device outlet and analyzed. Bright and fluorescence microscopy confimred effective mixing and depletion of blood cells.

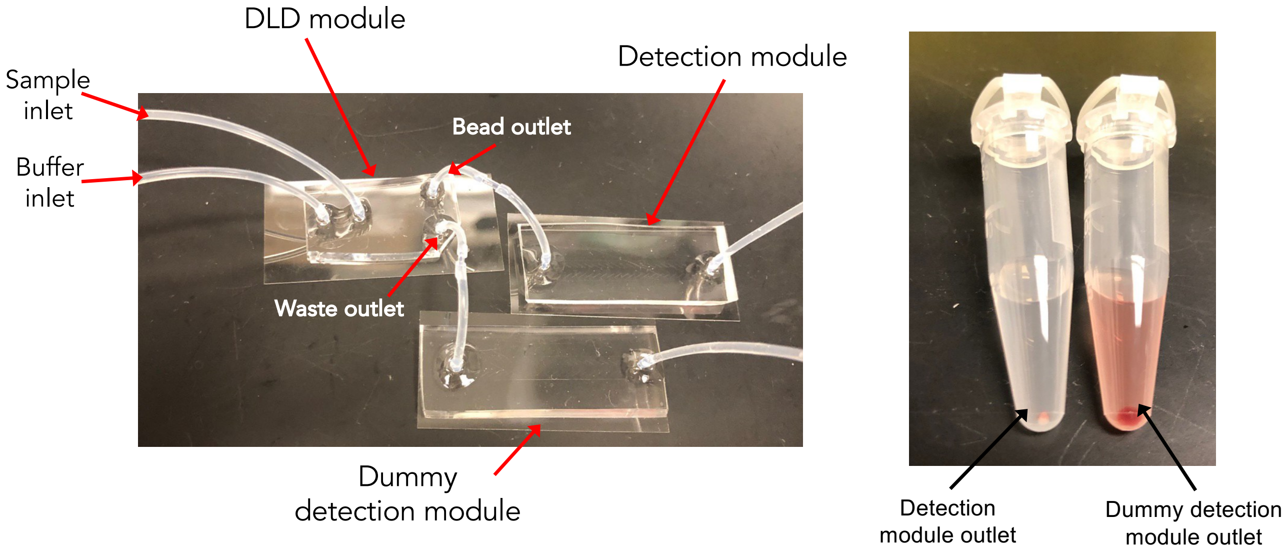


**Figure S4. Integration of the depletion and detection modules.** The depletion module was connected to primary and dummy detection modules. A blood sample spiked with 15-µm microbeads was injected into the sample inlet of the depletion module, while a sheath buffered solution was injected through the buffer inlet. After separation, beads and blood were collected from the primary and dummy detection modules, respectively. The image at right shows the effective separation of beads from blood cells.

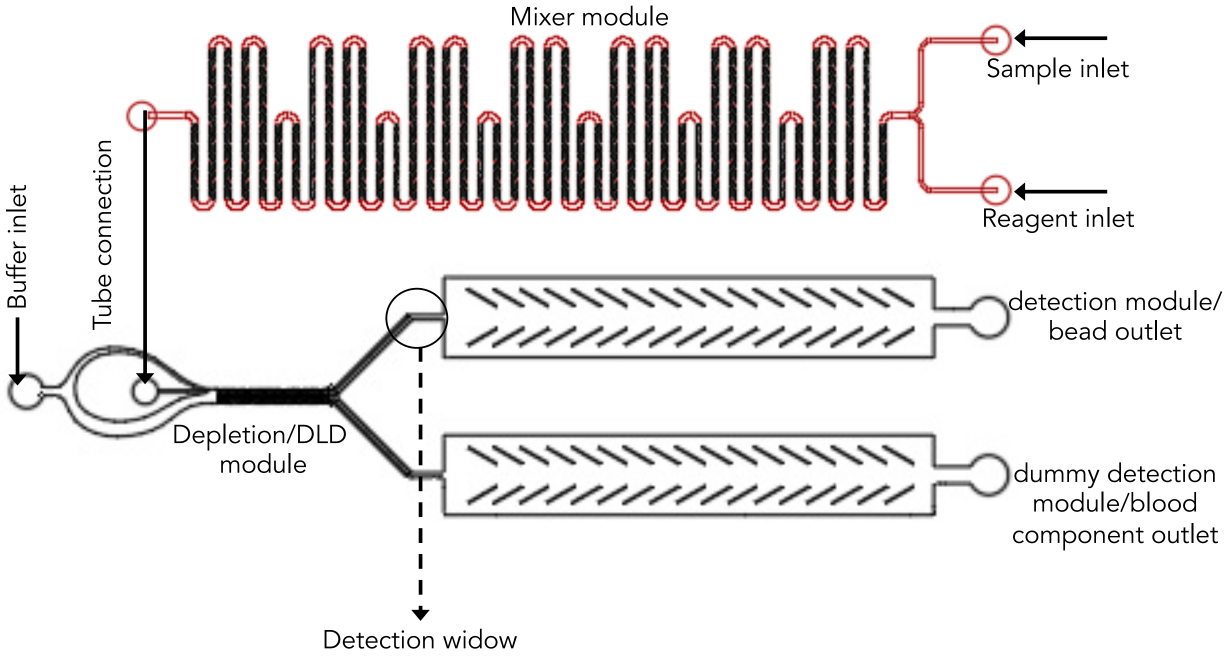


**Figure S5. Integrated RT-ELISA platform mask design.** The device mask was designed using AutoCAD software and then used to fabricate the silicon masters.

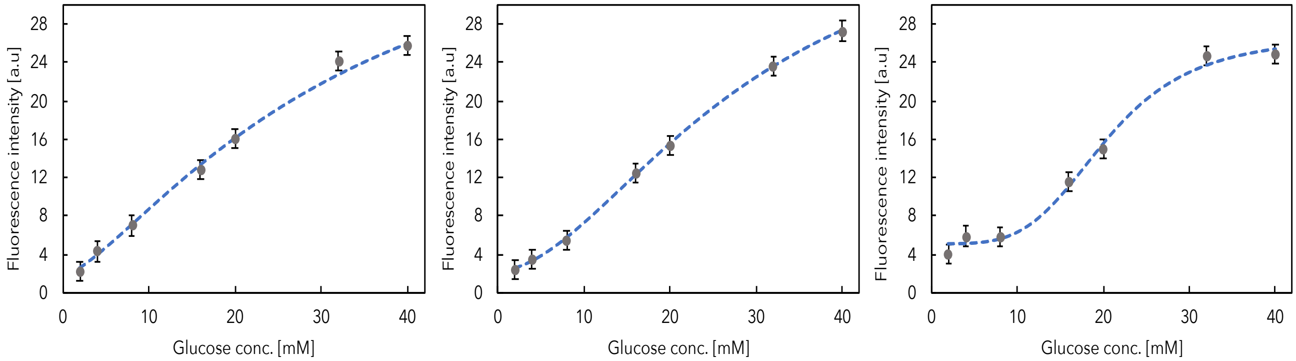


**Figure S6. Replicates for glucose standard curve derivation.** Human blood spiked with known concentrations of glucose was run through the RT-ELISA device. A red laser (642 nm/5 mW) was used to excite the Cy5 fluorophore on the glucose-bound beads, with signal intensity detected using a sCMOS camera. We employed a custom program to measure fluorescence intensity. These experiments were repeated three times at each concentration, with the intensity of ≥200 beads measured. Error bars show SD among beads. Endogenous glucose was measured using a conventional glucose meter and subtracted as baseline before calculating the spiked concentrations. Non-linear regression analysis is used for curve-fitting.

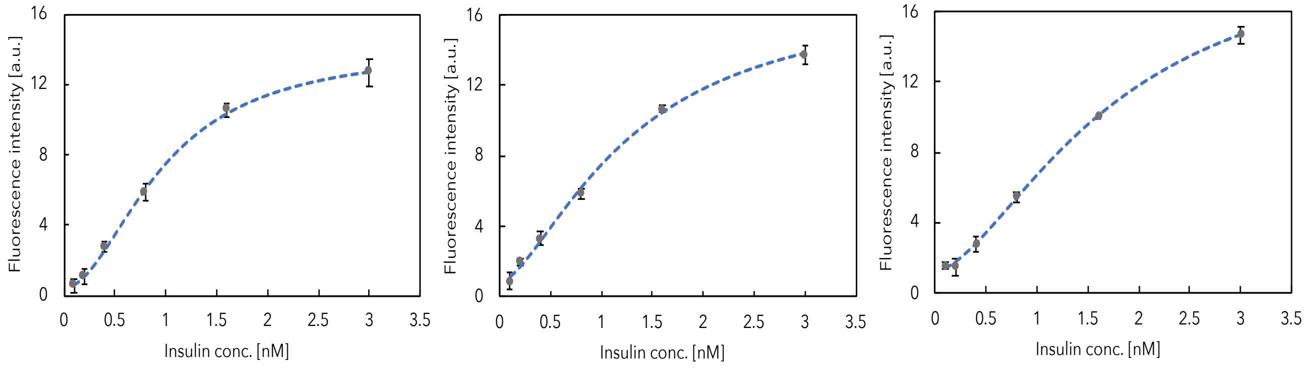


**Figure S7. Replicates for insulin standard curve derivation.** Human blood spiked with known concentrations of insulin was run through the RT-ELISA device. A green laser (520 nm/40 mW) was used to excite the R-PE detection fluorophore. Detection and analysis were performed as described in **Figure S6**. The fluorescence signal intensity from a blank blood sample was measured to assess endogenous insulin and was subtracted from the measured signal at each concentration. Non-linear regression analysis is used for curve-fitting.

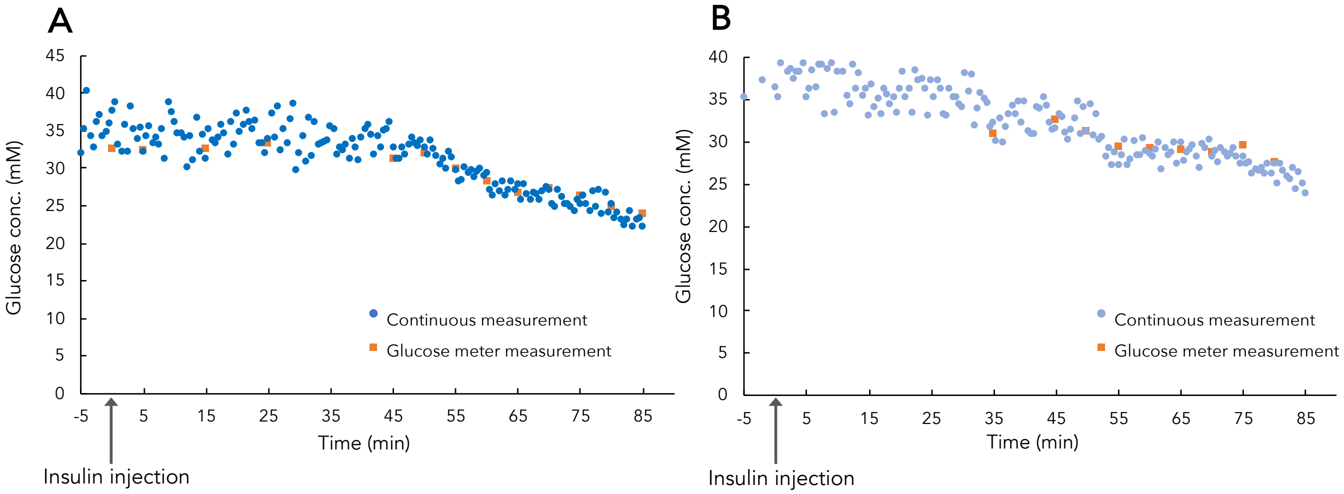


**Figure S8. Glucose monitoring data in live rats after injection of two different insulin formulations** (related to Figure 7). **A)** Humulin R injection and **B)** Humulin N injection. In **B,** for the first 35 mins, the blood glucose was greater than the detection limit of the handheld glucose monitor and could not be measured.

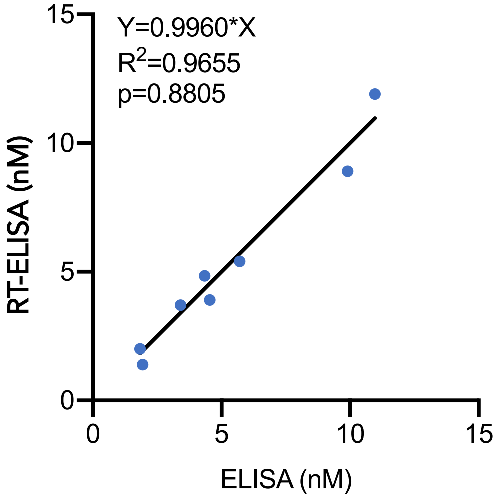


**Figure S9. Regression statistical comparison of RT-ELISA and ELISA for *in vitro* secretion assays**. RT-ELISA measurements for insulin plotted against standard ELISA measurements. Regressions were fitted using GraphPad Prism 8's straight line non-linear regression, with the y-intercept constrained to equal 0. Best fit for insulin measurements did not differ from the null hypothesis Y = X, suggesting that the measurements between methods are highly correlated.

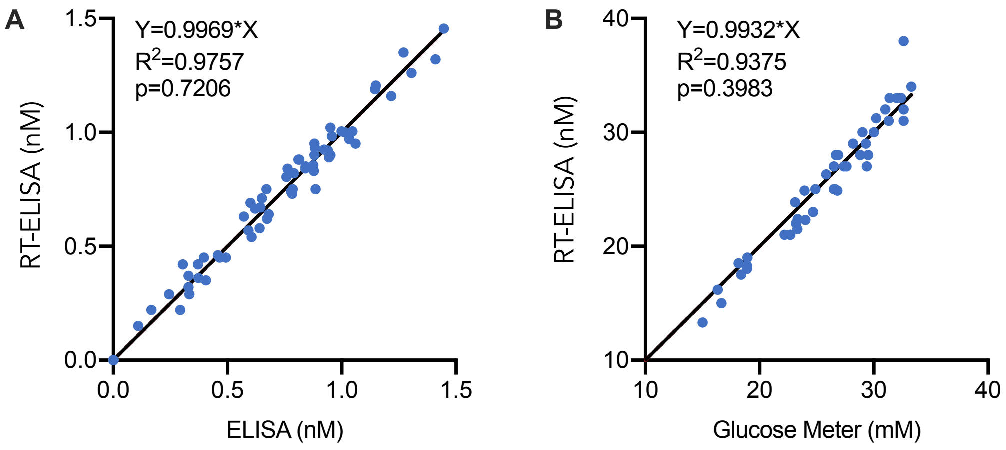


**Figure S10. Regression statistical comparison of RT-ELISA and ELISA or glucose monitor for *in vivo* rat experiments. (A)** RT-ELISA measurements for insulin plotted against standard ELISA measurements. (**B**) RT-ELISA measurements plotted against glucose meter readings. Regressions were fit using GraphPad Prism 8's straight line non-linear regression, with the y-intercept constrained to equal 0. The line of best fit for both insulin and glucose measurements were not different from the null hypothesis Y = X, suggesting that the measurements between methods are highly correlated.

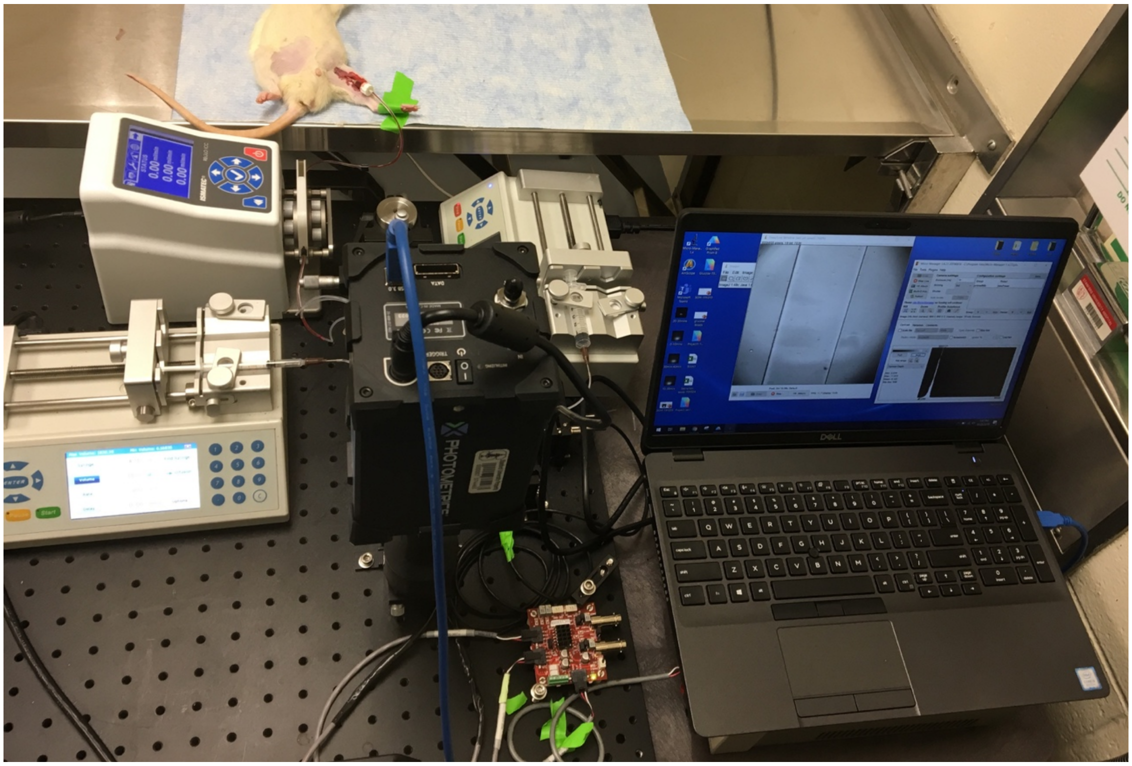

**Figure S11. Animal experiment setup.** A peristaltic pump was used to withdraw blood from rats at 15 μL/min, and two syringe pumps were used to inject the reaction solution and buffered solution required for depletion into the device. As beads pass through the detection window, their fluorescence intensity was measured using a sCMOS camera.

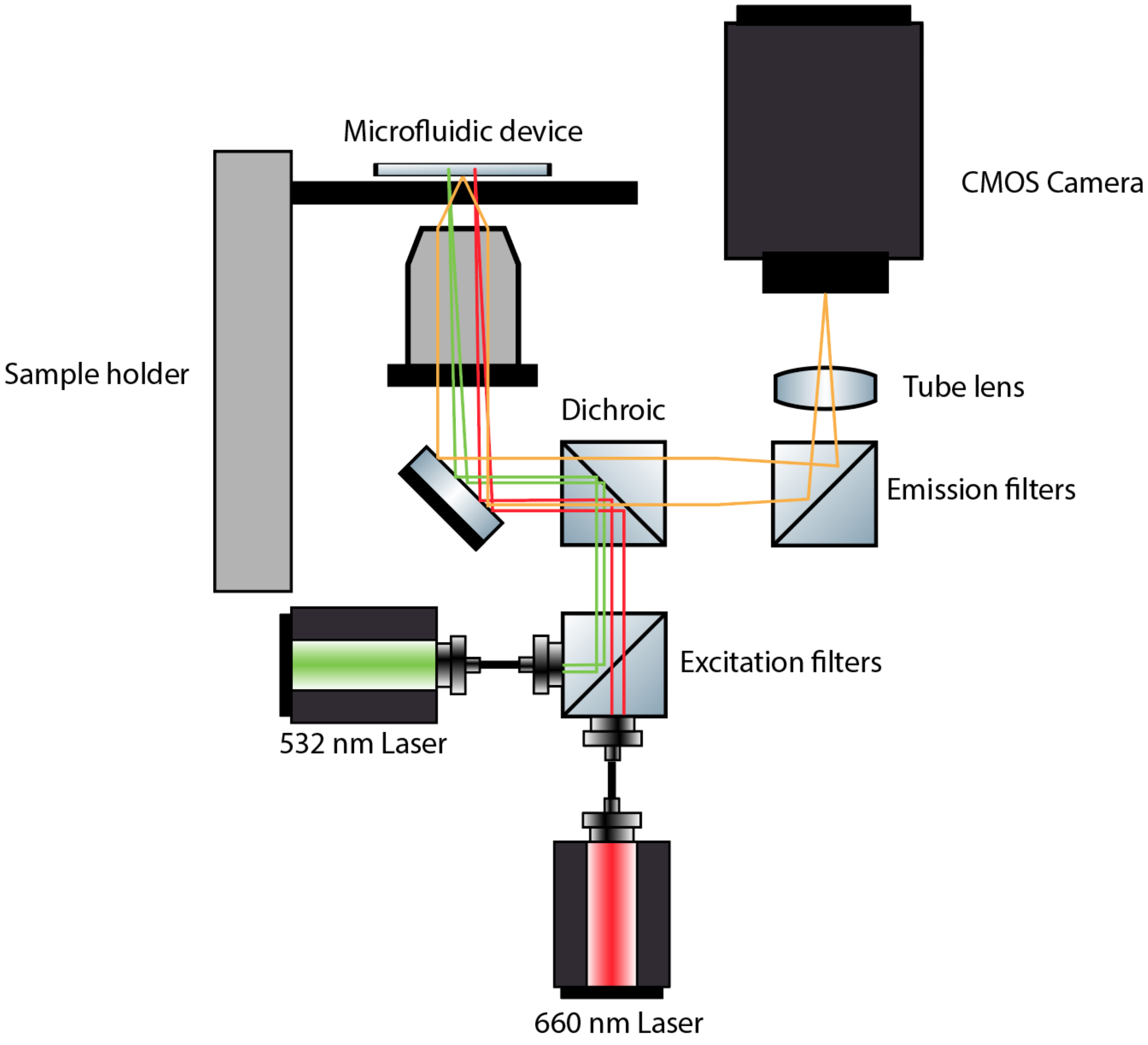


**Figure S12. Schematic of the imaging setup.**

Microscope videos

The following are descriptions of the videos, enclosed with this document:

1) **Video S1:** DLD separation; that supplements the cell trajectories depicted in Figure 3B of the main text. The video demonstrates the separation of fluorescently labeled microbeads from blood cells and free fluorescently tagged antibodies.
1) **Video S2:** Glucose and insulin beads passing through the detection window.
2) **Video S3**: Control; that shows glucose beads only fluoresce in their specific region on the upper part of detection window.
3) **Video S4**: Control; that shows insulin beads only fluoresce in their specific region on the bottom of detection window.
